# Protein stability and functional activity during nebulization: A comparative study of three nebulizer!

**DOI:** 10.1101/2020.05.09.085720

**Authors:** Niti Singh, Preeti Yadav, Prekcha Gaur, Manish Gaur, Awadh Bihari Yadav

## Abstract

Delivery of therapeutics protein to the lung offers effective treatments of lungs disease. Efficacy of delivered therapeutics molecules depends on integrity and stability of protein during nebulization. In this study, we compared three nebulizers: compressed air nebulizer (CAN), ultrasonic nebulizer (USN) and mesh nebulizer (MAN) to deliver aerosol dose, stability and functional activity of a model protein lysozyme. Lysozyme/BSA delivered dose assessed by indirect and direct method. It was shown CAN deliver 0.142±0.027 to 0.632± 0.09 ml of protein, USN deliver 0.511±0.119 to 1.688±0.173 ml and MAN deliver 0.238±0.006 to 0.731±0.013 ml of protein in the same time. Integrity of nebulized proteins were assessed by gel electrophoresis and circular diochorism. It was found integrity of lysozyme compromised in all three nebulizer maximum in CAN and minimum with MAN. The functional activity of protein was assessed after nebulization by turbidometry assay. The functional activity was compromised by all three nebulizer upto some extent. In conclusion, nebulization compromise protein stability: this impact depend on nebulization techniques as well as nature of protein. The CAN deliver protein more precisely in small amount in comparison to the other nebulizer.

Graphical Abstract

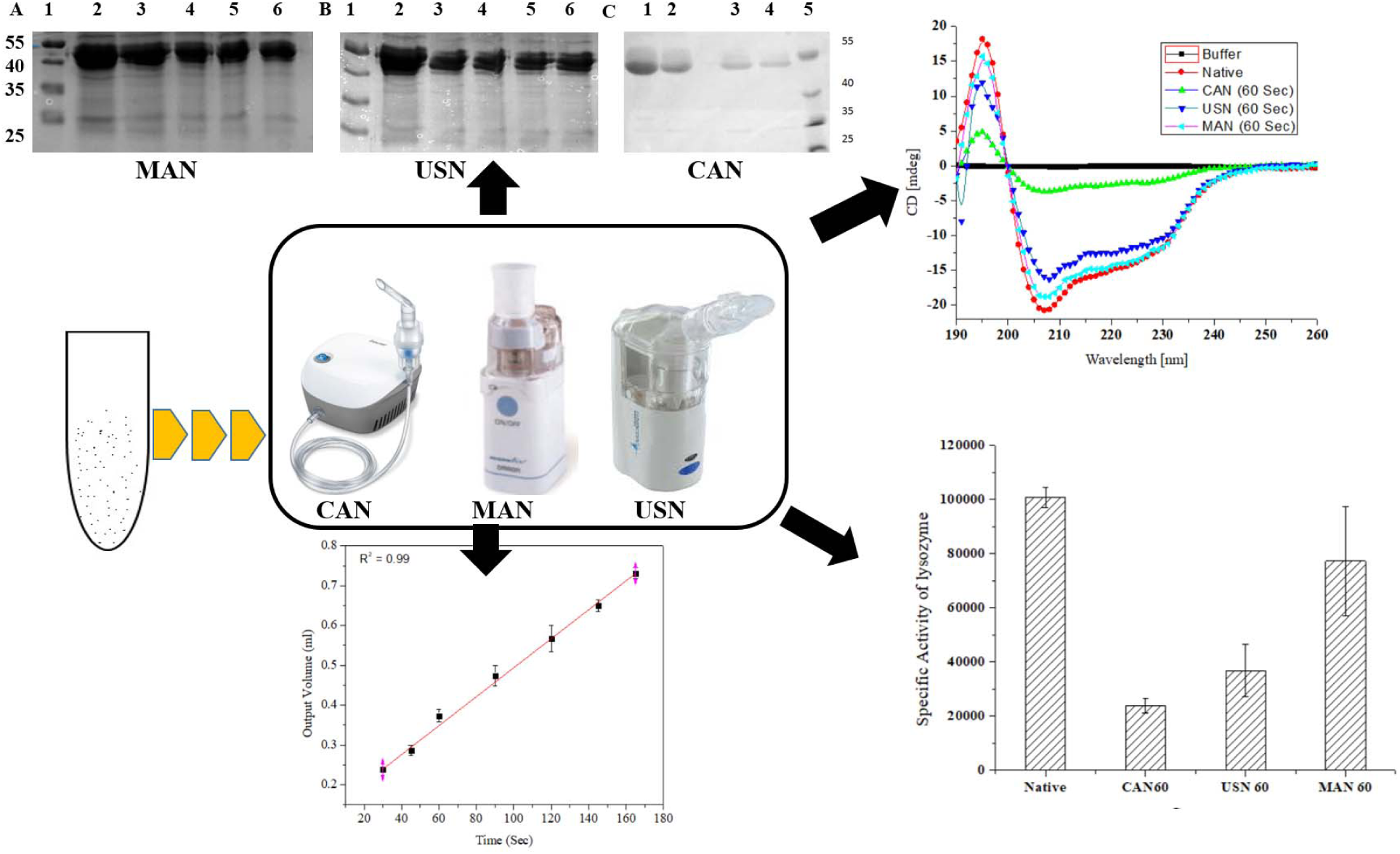

## Introduction

Lung diseases are the most common diseases, which includes asthma, chronic obstructive pulmonary disease (COPD), Cystis fibrosis, tuberculosis and lung cancer, spread worldwide [1]. Lung disease associated pathologies includes inflamed airways, allergies, infections, long term wet cough and other condition which ultimately compromise the healthy functioning of a lung [2]. Main causes of the lung disease are smoking, infections and pollution, which lead to the damage of the lung surface. According to WHO report, 235 million people is suffering from asthma worldwide and 30% of Indian population suffers from lung disease [3, 4]. At present, different treatments used for the treatment of Asthma & lung inflammation are delivered thorough the inhalation route [5]. Inhalation route offer advantages on conventional route and provide quick relief from the life threatening condition developed in asthma and other diseases by delivering quick therapeutics molecules on the lung surface [6, 7]. In asthma and lung inflammation steroid based therapeutics are in used which compromise immune system and add more side effect [8]. Protein/peptide based therapeutics molecules offers advantages over steroid based therapy to not interfere with host immune system, target specific activity and less side effect in comparison to other therapy [9, 10].

Therapy based on protein/peptide limited by their stability & integrity to deliver at the site of disease. Several therapeutic proteins/peptides under different developmental stage for the treatment of lungs disease by nebulization method [11]. The human deoxyribonuclease I and dornase-α are used for the treatment of cystic fibrosis and lung infections respectively [12, 13], nebulized recombinant secretory leukocyte protease inhibitor (rSLPI) show good activity in lung inflammation site and has been proposed for the treatment of asthma [14]. A. Artigas et.al. demonstrated that nebulized heparin control pulmonary inflammation through macrophage in rat lung Injury model [15]. Acute lung injury and pulmonary inflammation can be reduced by aerosolized indomethacin in to a rat animal model [16]. Other groups demonstrated that aerosolized pulmonary surfactant inhaling can reduce proinflamatory cytokines in bronchoalveolar lavage fluid (BALF) [17]. Y. Chen et.al. reported that nebulized mycobacterium vaccine can protect against asthma in mice [18]. Nebulized anticoagulants can also attenuate lung inflammation [19].

Efficacy of therapeutics molecules depends on many factors, such as stability, integrity of molecules during and after nebulization. High-frequency acoustic waves based nebulizer delivered 70% size range of peptide for deep lung delivery without any compromise in their integrity [20]. A nebulizer based on acoustomicrofluidic capable to nebulized epidermal growth factor receptor (EGFR) into fine aerosol of mass median aerodynamic diameter of approximately 1.1 μm with no significant degradation observed after nebulization [21]. In inhaled therapy it’s very critical and essential to preserve stability and integrity of formulation during nebulization [22].

The aim of this study was try to find out which nebulizer able to deliver protein/peptide without or minimum compromise their stability during nebulization. At present nebulizer are used based on three principles: compressed air nebulizer, ultrasonic vibration & mesh nebulizer. Furthermore all three nebulizer was analyzed for dose delivered as output dose, stability, integrity and functional activity of nebulized protein.

## 2. Materials and Methods

### 2.1. Materials

Bovine Serum albumin (BSA), Lysozyme, Micrococcus luteus, Tween-20 and all electrophoresis reagents from from Sigma, Banglore, India, Phosphate Buffer Saline (PBS), Coomassie Brilliant Blue G-250 dye, Alkaline phosphate buffer (pH 7.4), TrisHCl buffer, Acetic acid and Ethanol from Hi Media, India, Molecular Weight marker 10-250 kDa (Fermentas) and Glycerol, Methanol (Molychem).

### 2.2. Methods

#### 2.2.1. Nebulization and metered dose delivery

In this study we used three different nebulizers: Compressor air nebulizer (CAN), ultrasonic nebulizer (USN) and mesh nebulizer (MAN) to deliver lysozyme or BSA as model molecules.

CAN (NE-C28, Omron Healthcare Co., Ltd) works between the temperatures range 10 ºC – 40 ºC and is capable of generating a particle size of ∼3µm, with aerosol output rate of 0.06 ml/min, and a nebulization rate of 0.4 ml/min with sample reservoir between 2–7 ml. The Mass Median Aerodynamic Diameter (MMAD) of generated particles was 3 µm.

USN (BD5200, BremedTodi (PG)-Italy) works between the temperature range 10 ºC – 40 ºC. The sample reservoir volume for this nebulizer is 3 ml and the average nebulization rate is 1 ml/min. The particles generated by ultrasonic nebulizer fall between size ranges of 0.5 to 4 µm.

Mess nebulizer (NE-U22, Omron Healthcare Co., Ltd) contains mesh/membrane with 1000-7000 laser drilled holes, which vibrates at the top of the sample reservoir due which a pressures out a mist of very fine droplets through the holes. The sample reservoir volume between 3ml-5ml. The aerosol output rate is approx. 0.85 ml/min. The average nebulizer flow rate is 0.429ml/min.

Metered dose delivered by the three different nebulizers at different time points was analyzed by indirect and direct method.

In the indirect method, the dose delivered was determined by subtracting final volume remain in reservoir with the initial sample volume after nebulization. In brief, 3 ml of sample was taken into CAN, USN or MAN. Sample was nebulized for different time points: 30, 45, 60, 90, 120, 145 and 165 sec. After nebulization, delivered dose was calculated by subtracting the remaining volume of sample from initial volume of sample in reservoir. The same experiment was repeated for five times (n=5).

In the direct method, the dose delivered was determined by measuring total protein in the collected volume after different time nebulization by measuring collected sample in 50 ml falcon tube or measuring protein in collected sample by Bradford or Lorry method. In brief, 3 ml of protein sample was taken into nebulizer and nebulized for different time point 30, 45, 60, 90, 120, 145 & 165. 30 µl of the nebulized was diluted to make 500 µl in buffer and equal amount of Bradford reagent was added to quantified protein by measuring absorbance at 595nm by UV-Visible spectrophotometer (Evolution 201, Thermo). The same experiment was repeated for three times (n=3)

#### 2.2.2. Stability of Nebulized Protein

##### 2.2.2.1. Gel Electrophoresis

Protein integrity after nebulization by CAN, USN or MAN nebulizers were studied by gel electrophoresis (10% SDS PAGE). In brief, nebulized protein aerosols were collected at different time point and collected samples were loaded on poly acrylamide gel along with appropriate molecular weight marker. The electrophoresis was performed at standard condition and gel was stained with Coomassie Brilliant Blue R-250 dye.

##### 2.2.2.2. Circular Diochorism (CD) Spectroscopy

Stability of lysozyme was studied after nebulization by CAN, USN & MAN by circular diochorism (CD) spectroscopy. In brief nebulized lysozyme sample was collected at 60 sec. These collected sample was analyzed for their CD spectra between wavelength 190-270 nm in the equal amount of protein sample. This experiment was performed in triplicate to record CD spectra. Water and Buffer was taken as blank for this experiments.

#### 2.2.3. Lysozyme activity by turbidity assay

The functional activity of lysozyme was assessed by measuring turbidity at different time points 30, 45, 60, 90, 120, 145 & 165 sec of nebulized sample by three different nebulizer [23]. In brief, Lysozyme solution was prepared into 66 mM potassium phosphate buffer (pH 6.24) and micrococcus luteus (0.015% (w/v)) solution into the same buffer. 2.5 ml of bacterial solution were transferred into a quartz cuvette and reference absorbance was read at 450 nm. 100 µl of lysozyme solution were added and mixed quickly and absorbance was recorded for 5 min at 25 ºC and each reaction was performed in triplicate. Enzyme activity into different sample was calculated by following equation mentioned below:

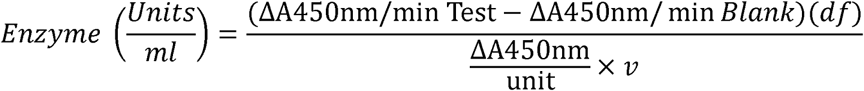

Where df = Dilution factor

ΔA450 nm/unit = Change in absorbance at A450nm/ per Unit of enzyme

v = Volume of enzyme (in milliliter) used in assay

#### 2.2.5. Statistical analysis of the results

The significance levels of the results between groups were calculated by one-way Analysis of variance (ANOVA) which including post hoc testing by Tukey to calculate individual p-values. The symbols and significance levels displayed are listed in Table 1.

**Table 1.** Significance levels and symbols to indicate them throughout this study.

| Symbol | Meaning |
| --- | --- |
| Ns | $P > 0.05$ |
| * | $P \leq 0.05$ |
| ** | $P \leq 0.01$ |
| *** | $P \leq 0.001$ |

## 3. Results and Discussion

### 3.1. Metered dose delivered by different nebulizer

The focus of this study was to establish dose of protein delivered by different nebulizers and to find out whether integrity of delivered protein was compromised during nebulization, which ultimately affect efficacy of protein. Therefore protein sample was nebulized for same time using all three nebulizer CAN, USN & MAN. We use phosphate buffer for lysozyme study. A statistically significant data using statistical design to represented with sufficient number of replicate of experiments (Table 1).

The output volume of lysozyme/BSA was assessed by measuring nebulized volume of sample by indirect method. We observed correlation of output volume nebulized between 30 sec to 165 sec by CAN was R^2^= 0.99, for USN R^2^ = 0.99 and for MAN R^2^ = 0.99 (Fig. 1A). We also assessed output dose collected into 50 ml falcon tube by direct method at different time point between 30 sec to 165 sec. Protein in collected sample was quantified by Lowry method. The correlation of output dose was found to be for CAN R^2^ = 0.98, USN R^2^ = 0.99 and MAN R^2^ = 0.99 (Fig. 2A).

**Fig. 1.**
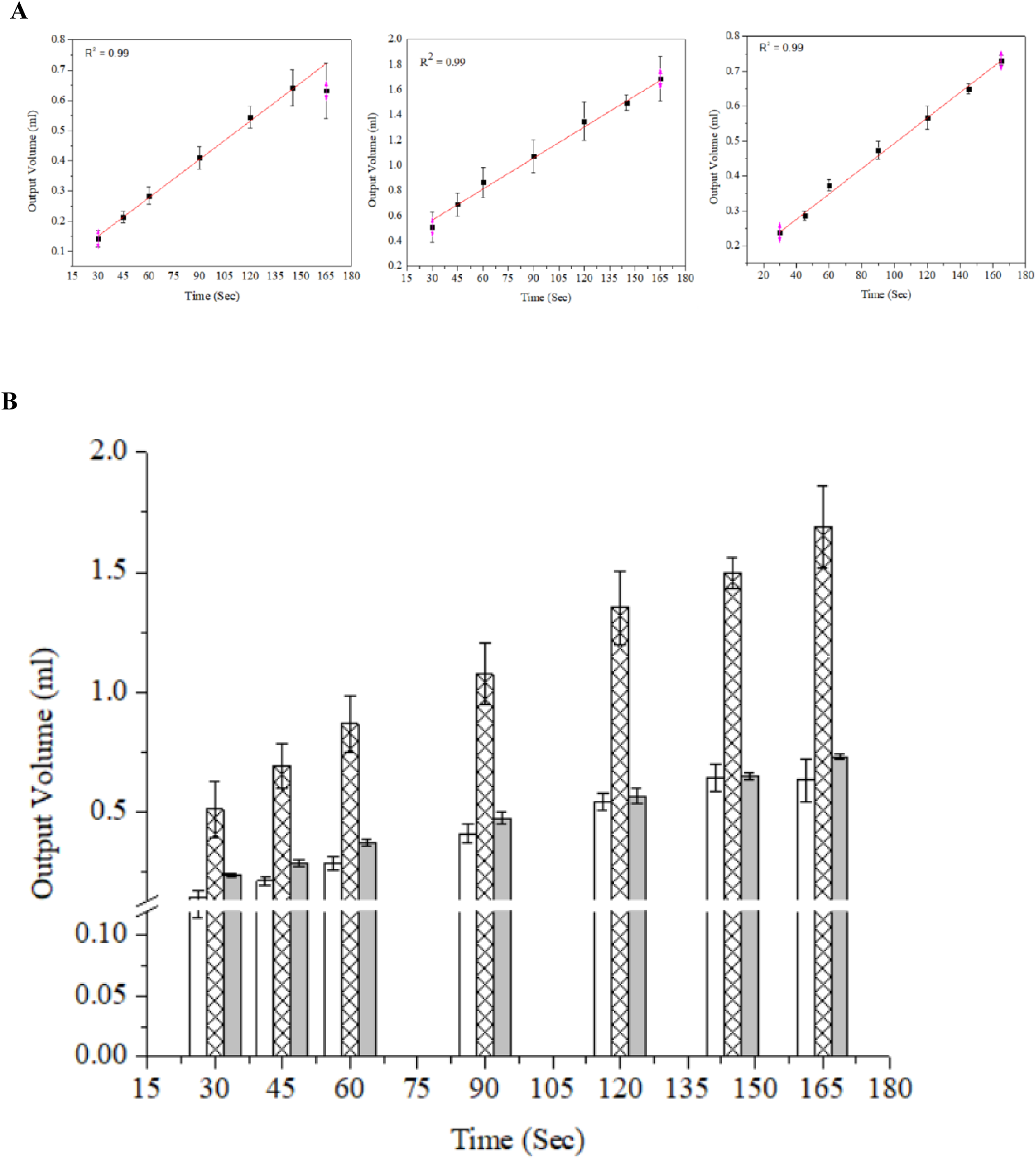
**(A)** Correlation of output volume with reference to time of nebulized protein by CAN (R^2^ = 0.99), USN (R^2^ = 0.99) and MAN (R^2^ = 0.99).**(B)** Dose delivered by CAN (hollow bar), MSN (cross line bar) and MAN (light gray bar) after nebulization and analyzed by indirect method, n=5 (mean ± SD).

**Fig. 2.**
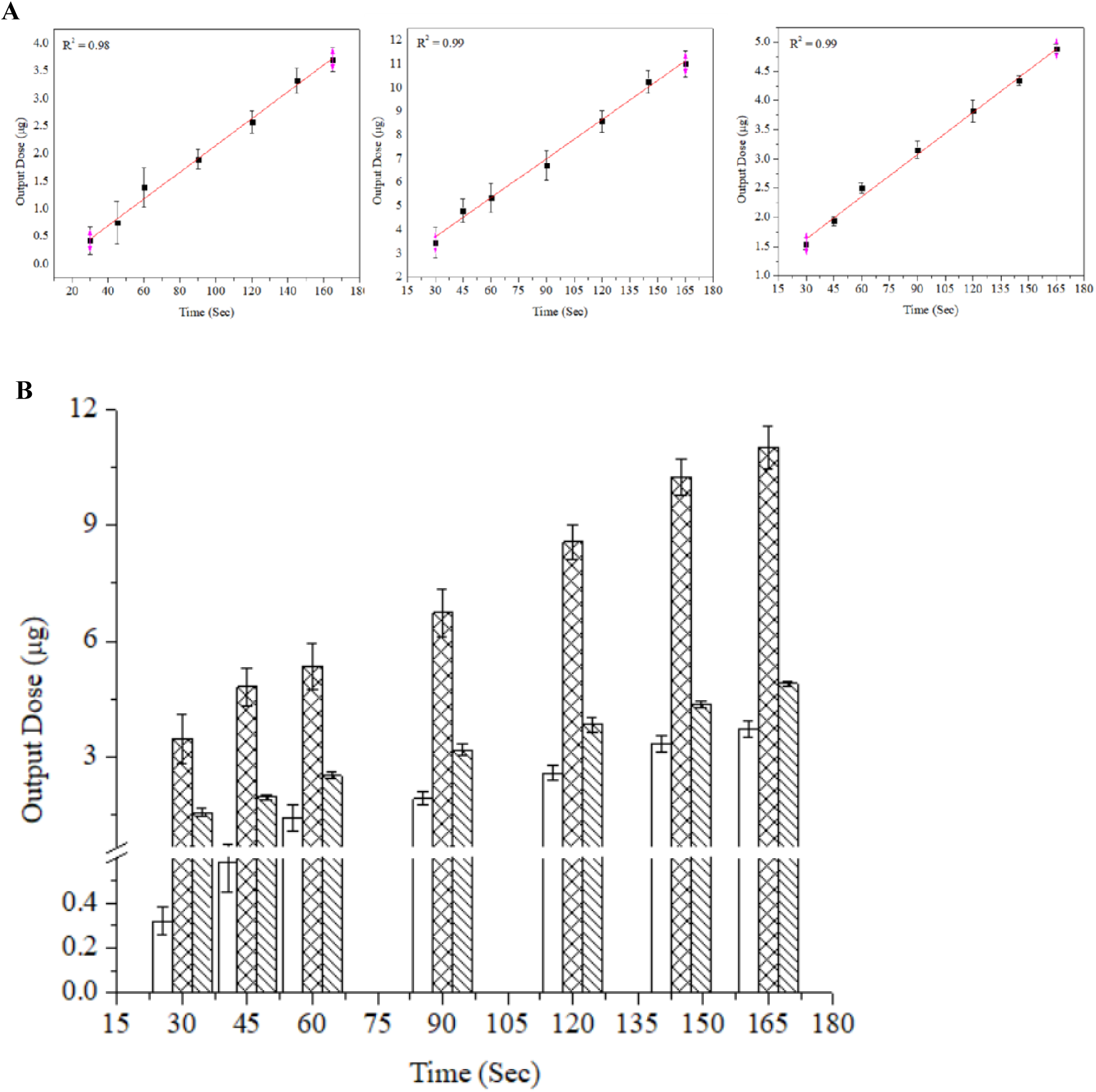
**(A)** Correlation of output dose with reference to time of nebulized protein by CAN (R^2^ = 0.98), USN (R^2^ = 0.99) and MAN (R^2^ = 0.99). **(B)** Dose delivered by CAN (hollow bar), MSN (cross bar) and MAN (cross section bar) diagram at different time point by direct method, n=5 (mean ± SD).

The dose delivered assessed by indirect method was 0.142±0.027 to 0.632± 0.09 ml CAN, 0.511±0.119 to 1.688±0.173 ml USN and 0.238±0.006 to 0.731±0.013 ml MAN 30 to 165 sec (Fig.1 B). The dose delivered also assessed by direct method was 0.318±0.061 to 3.71±0.215 µg CAN, 3.45±0.653 to 11.02±0.553 µg USN and 1.55±0.098 to 4.88±0.079 µg in 30 to 165 sec (Fig.2 B). The compressed air nebulizer can deliver small dose of protein more precisely in comparison to USN and MAN. The dose delivered by different nebulizer was also assessed by direct method. Protein was estimated in sample collected at different time point after nebulization by CAN, USN & MAN nebulizer by Bradford/lowery method. In this study it was shown USN and MAN can deliver large amount of protein in small time in comparison to CAN.

### Stability of protein

The impact of nebulization on protein integrity and stability was studied by gel electrophoresis and CD spectroscopy of sample nebulized by CAN, USN & MAN nebulizer. This experiment were performed with a protein solution into a recommended strength and pH of buffer. Stability of nebulized proteins is very crucial for the efficacy of protein. Degradation of protein was observed in SDS PAGE following CAN, USN & MAN nebulization (Fig. 3).

**Fig. 3.**
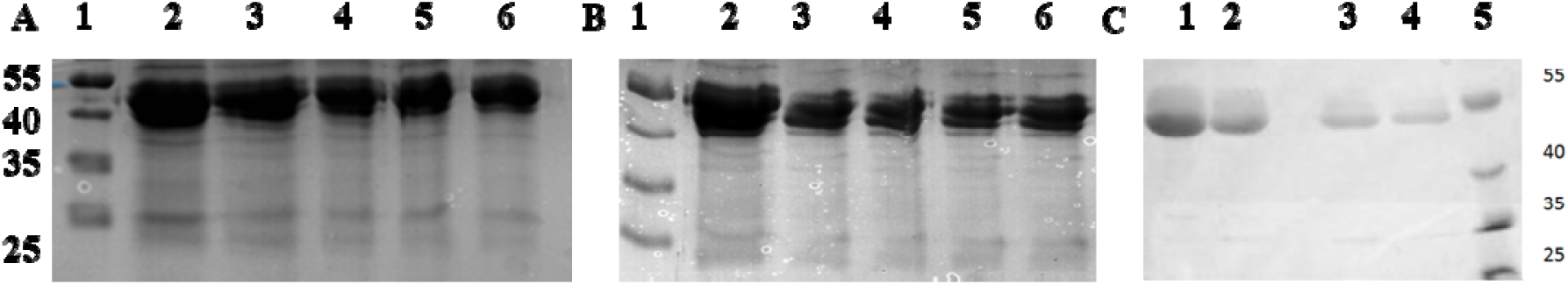
**SDS PAGE analysis of sample after nebulization for different time point by (A)** MAN lane 1 molecular weight marker (fermentas, SM1811), lane 2 native protein, lane 3-6 protein in different time interval (45 sec, 120 sec 165sec, 195 sec), **(B)** by UAN lane 1 molecular wieight marker, lane 2 native protein lane 3-6 protein in different time interval (45 sec, 120 sec 165sec, 195 sec) **(C)** by CAN lane 5 molecular wieight marker, Lane 4 native protein Lane 3, 2 and 1 protein sample nebulized for different time point (5, 7 & 9 min).

Near UV CD spectra provide information about the environment around aromatic group in a protein. Difference in the spectra (peak shifting) of a protein indicate that there is change into surrounding environment around aromatic group in a protein. Near UV CD spectra of lysozyme was assessed after nebulization of CAN, USN & MAN (Fig. 4). Buffer and protein solution was used as reference sample. The sample nebulized by different nebulizer for 60 sec was showing different amount of tertiary structure compromization: maximum compromised by CAN, USN and minimal structure compromised by MAN.

**Fig. 4.**
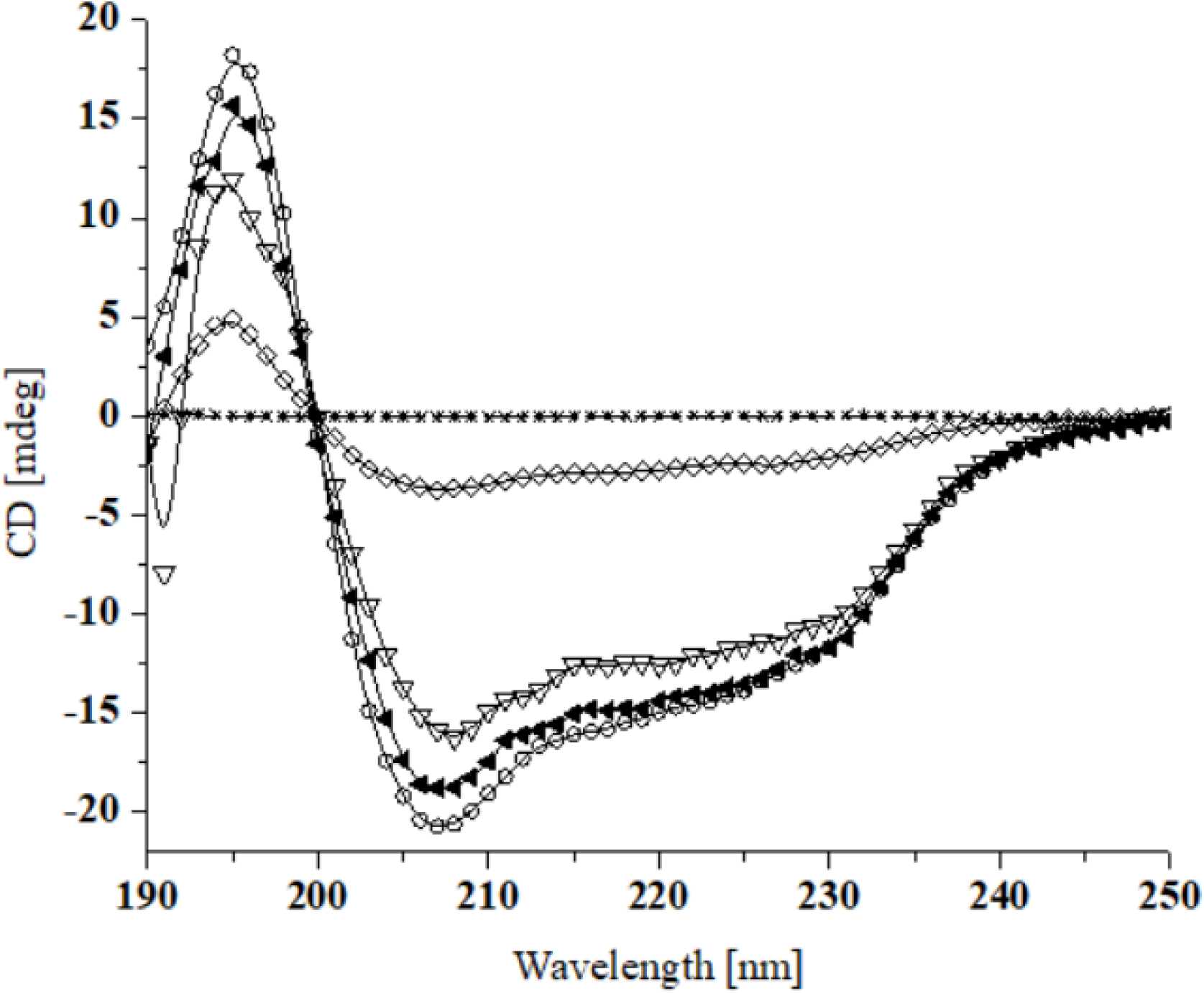
CD spectra of nebulized buffer (-*-), CAN (-⍰-), USN (-Δ-), MAN (-▴-) and Native (-○-) of lysozyme sample for 60 sec and spectra recorded at near UV region 190-270 nm of 100 ug of protein in each sample n = 3 (mean±SD).

### Activity of Lysozyme

The integrity and stability compromised during nebulization whether it affect lysozyme activity was investigated by turbidity assay. Lysozyme sample nebulized by CAN, USN & MAN nebulizer for 60 sec. The lysozyme activity of nebulized sample were compared with native lysozyme solution without any nebulization (Fig. 5). Functional activity of lysozyme was also assessed in the sample after nebulization for different time (Data not shown).

**Fig. 5.**
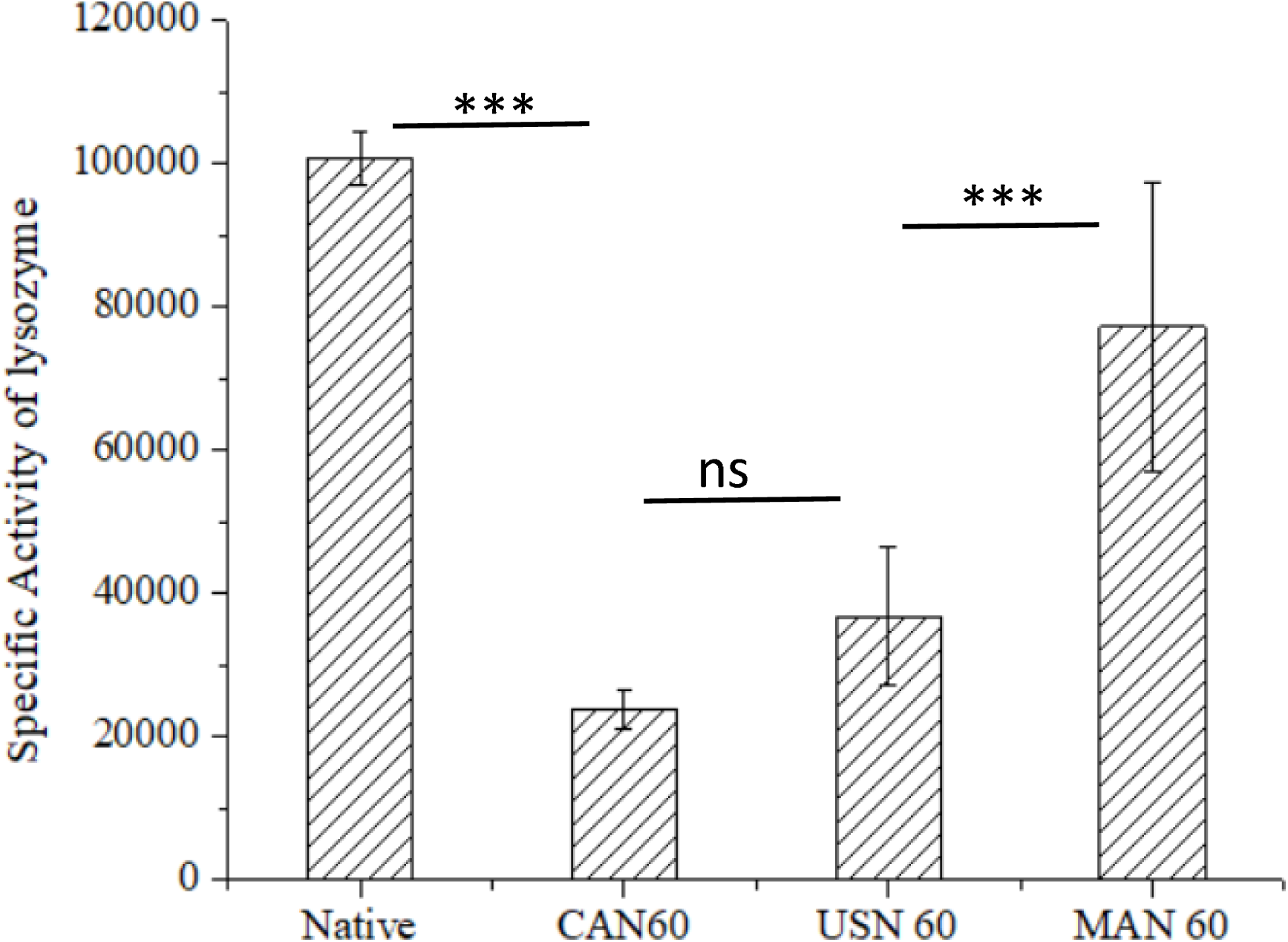
Lysozyme activity in nebulized aerosol sample collected for 60 sec by CAN, USN or MAN nebulizer by turbidometry assay, n = 5 (mean±SD).

During nebulization by CAN for 60 sec 82.6% lysozyme activity was lost, nebulization by USN 74.4% and nebulization by MAN 33.2% lysozyme activity was lost. There are different factor which attribute to the activity loss of lysozyme sample which includes heating, aggregation of protein and liquid air interface formation [24]. In CD spectra it was shown maximum structural loss was in sample nebulized by CAN (Fig.4). Activity loss of lysozyme during nebulization can be minimized by reducing temperature of sample reservoir during nebulization and to reduce interfacial stress by adding suitable excipient [24]. An appropriate combination excipient and temperature control required to achieve optimal protection and retain optimal functional activity of lysozyme during nebulization [25, 26].

## 4. Discussion

We studied the dose delivered by three different nebulizer in the same time and how it affect stability and activity of a protein during nebulization. It was found that dose delivered by compressed air nebulizer more precise in small amount in the same time dose delivered by ultrasonic and mesh nebulizer. All three nebulizer affect stability of protein sample during nebulization: maximum compromised by CAN, USN and minimum by MAN. These stability compromised due to many factor such as heat generation, air water interface formation & fragmentation of therapeutic molecules during nebulization by different nebulizer. In 2015 W. Friess *et. al*. explain protein instability due to unfolding & aggregation at the air-liquid interface [11]. They also demonstrated stability for protein vary and depend on the nature protein [27]. Stability of protein sample can be improved by minimizing heat generation during nebulization. It can be also improved by incorporating suitable excipient to protect samples exposure to air and surface interface which also responsible for denaturation of protein sample during nebulization. It observed that Tertiary structure of protein affected by nebulization. Different nebulizer affect different level of tertiary structure of a protein. Slight change in tertiary structure significantly affect activity of protein.

Protein stability compromised during nebulization is a bottle neck to use of this convenient administration method to deliver protein formulation into the lung for the treatments of many disease such as tuberculosis, lung inflammation, COPD, cystic fibrosis. During nebulization it was reported that formation of aggregates and degradation of protein affect efficacy of therapeutic protein [24, 28]. Ultrasonic nebulizer also affect protein efficacy around 50% of activity loss was reported during nebulization of protein solution [29].

Conclusion of our study (i) compressed air nebulizer deliver small dose more precisely (ii) ultrasonic and mesh nebulizer delivered high amount in comparison to CAN. Protein nebulized by CAN, USN & MAN nebulizers is prone for degradation. Protein nebulized by CAN show high degradation in comparison to the USN & MAN. Maximum activity loss of lysozyme observed in compressed air nebulized sample, no or very less activity loss was observed in sample nebulized by mesh nebulizer.

## Author contributions

### Funding Source

This work was supported by the **Core Grant CRG/2018/002135** grant by Science and Engineering Research Board (SERB), DST, Government of India.

## Acknowledgment

We are thankful to Science and Engineering Research Board (SERB) New Delhi, India for providing financial support for this project.

